# LazySampling and LinearSampling: Fast Stochastic Sampling of RNA Secondary Structure with Applications to SARS-CoV-2

**DOI:** 10.1101/2020.12.29.424617

**Authors:** He Zhang, Liang Zhang, Sizhen Li, David H. Mathews, Liang Huang

**Affiliations:** Baidu Research USA, Sunnyvale, CA; School of Electrical Engineering & Computer Science, Oregon State University, Corvallis, OR; Dept. of Biochemistry & Biophysics, University of Rochester Medical Center, Rochester, NY 14642, USA; Center for RNA Biology, University of Rochester Medical Center, Rochester, NY 14642, USA; Dept. of Biostatistics & Computational Biology, University of Rochester Medical Center, Rochester, NY 14642, USA

## Abstract

Many RNAs fold into multiple structures at equilibrium. The classical stochastic sampling algorithm can sample secondary structures according to their probabilities in the Boltzmann ensemble, and is widely used. However, this algorithm, consisting of a bottom-up partition function phase followed by a top-down sampling phase, suffers from three limitations: (a) the formulation and implementation of the sampling phase are unnecessarily complicated; (b) the sampling phase repeatedly recalculates many redundant recursions already done during the partition function phase; (c) the partition function runtime scales cubically with the sequence length. These issues prevent stochastic sampling from being used for very long RNAs such as the full genomes of SARS-CoV-2. To address these problems, we first adopt a hypergraph framework under which the sampling algorithm can be greatly simplified. We then present three sampling algorithms under this framework, among which the LazySampling algorithm is the fastest by eliminating redundant work in the sampling phase via on-demand caching. Based on LazySampling, we further replace the cubic-time partition function by a linear-time approximate one, and derive LinearSampling, an end-to-end linear-time sampling algorithm that is orders of magnitude faster than the standard one. For instance, LinearSampling is 176× faster (38.9s vs. 1.9h) than Vienna RNAsubopt on the full genome of Ebola virus (18,959 *nt*). More importantly, LinearSampling is the first RNA structure sampling algorithm to scale up to the full-genome of SARS-CoV-2 without local window constraints, taking only 69.2 seconds on its reference sequence (29,903 *nt*). The resulting sample correlates well with the experimentally-guided structures. On the SARS-CoV-2 genome, LinearSampling finds 23 regions of 15 *nt* with high accessibilities, which are potential targets for COVID-19 diagnostics and drug design.

See code: https://github.com/LinearFold/LinearSampling

## 1. Introduction

RNAs are involved in many cellular processes, including expressing genes, guiding RNA modification (1), catalyzing reactions (2) and regulating diseases (3). Many functions of RNAs are highly related to their secondary structures. However, determining the structures using experimental methods, such as X-ray crystallography (4), Nuclear Magnetic Resonance (NMR) (5), or cryo-electron microscopy (6), are expensive, slow and difficult. Therefore, being able to rapidly and accurately predict RNA secondary structures is desired.

Commonly, the minimum free energy (MFE) structure is predicted (7; 8), but these methods do not capture the fact that multiple conformations exist at equilibrium, especially for mRNAs (9–12). To address this, McCaskill (13) pioneered the partition function-based methods, which account for the ensemble of all possible structures. The partition function can estimate the *base pairing probabilities p_i,j_* (nucleotide *i* paired with *j*), and the *unpaired probabilities q_i_* (*i* is unpaired).

However, the estimated base-pairing and unpaired probabilities *p_i,j_*’s and *q_i_*’s, being marginal probabilities summed over all possible structures, can only provide *summaries* of the exponentially large ensemble, but not direct and intuitive descriptions (14) which are needed in many scenarios. First, we often prefer to see a sample of representative structures according to their Boltzmann probabilities, which is more informative than the marginal probabilities (15). For example, we can use a set of samples to estimate the end-to-end distance of an RNA (12). Second, more importantly, we often want to predict the probability that a region is completely unpaired, known as the *accessibility* of that region, which plays an important role in siRNA sequence design (10; 11; 16; 17). Accessibility *cannot* be simply computed as the product of the unpaired probabilities for each base in the region because those probabilities are *not* independent.

To alleviate these issues, Ding and Lawrence (14) pioneered the widely-used technique of stochastic sampling, which samples secondary structures according to their probabilities in the ensemble. Their algorithm consists of two phases: the first phase computes the partition function (but not the marginal probabilities) in a standard bottom-up fashion, and the second “sampling” phase generates structures in a top-down iterative refinement fashion. This algorithm can estimate ensembles of structures, and predict the accessibility by sampling *k* structures and counting how many of them have the region of interest completely unpaired. Two popular RNA folding packages, RNAstructure (18) and Vienna RNAfold (19), both implement this algorithm.

However, widely-used as it is, the standard Ding and Lawrence sampling algorithm suffers from three limitations. First, its formulation and implementation are unnecessarily complicated (see Sec. B.1 for details). Secondly, the sampling phase repeatedly recomputes many recursions already performed during the partition function phase, wasting a substantial amount of time especially for large sample sizes.^1^ Finally, it relies on the standard partition function calculation that takes *O*(*n*^3^)-runtime, where *n* is the sequence length. This slowness prevents it from being used for long sequences including the full-genome of SARS-CoV-2.

To alleviate these three issues, we present one solution to each of them. We adopt the hypergraph framework (21; 22), under which the sampling algorithm can be greatly simplified. This framework conveniently formulates the search space of RNA folding, and then sampling can be simplified as recursive stochastic backtracing in the hypergraph.

Under this framework, we present three sampling algorithms. The first one, *Non-Saving Sampling*, is similar to, but much simpler and cleaner than, Ding and Lawrence’s, while the other two are completely novel. The second algorithm, *Full-Saving Sampling*, eliminates all redundant calculations by saving all computations from the partition-function phase. The third one, *LazySampling*, runs the fastest by only saving computations that are needed during the sampling phase, and is thus a clever trade-off between the first two.

Finally, we further improve LazySampling by replacing its *O*(*n*^3^)-time partition function calculation by our recently proposed *O*(*n*)-time approximate algorithm, LinearPartition (23). This combination gives rise to *LinearSampling*, an end-to-end linear-time sampling algorithm that is orders of magnitude faster than the standard algorithm. LinearSampling achieves 176× speedup (38.9s vs. 1.9h) compared to RNAsubopt on the full genome of the Ebola virus (18,959 *nt*).

More importantly, as the COVID-19 pandemic continues, it is of great value to find the regions with high accessibilities in SARS-CoV-2, which can be potentially used for diagnostics and drug design. However, previously there was no tool that can fast sample structures and calculate the accessibilities on such long sequences and consider global, long-range base pairs. LinearSampling is the first sampling algorithm to scale up to the whole SARS-CoV-2 genomes (~30,000 *nt*) without local window constraints, and can sample 10,000 structures in only 69.2 seconds. LinearSampling-derived accessibilities correlate well with the experimentally-guided structures (24), resulting in 23 regions of 15 *nt* with high accessibilities, which are potential targets for COVID-19 diagnostics and drug design.

## 2. Sampling Algorithms

We first formulate (in Sec.A) the search space of RNA folding using the framework of (directed) hypergraphs (21; 22) which have been used for both the closely related problem of context-free parsing (25)and RNA folding itself (22; 26). This formulation makes it possible to present the various sampling algorithms succinctly (see Sec. B), where sampling can be done in a top-down way that is symmetric to the bottom-up partition function computation. Finally, we present (in Sec. C)our LinearSampling algorithm which is the first sampling algorithm to run in end-to-end linear-time.

### A. Hypergraph Framework

For an input RNA **x** = *x*_1_…*x_n_*, we formalize its search space as a **hypergraph** 〈*V*(**x**), *E*(**x**)〉. Each **node** *v* ∈ *V*(**x**) corresponds to a subproblem in the search space, such as a span **x**_*i,j*_. Each **hyperedge** *e* ∈ *H*(**x**) is a pair 〈*node*(*e*), *subs*(*e*)〉 which denotes a decomposition of *node*(*e*) into a list of children nodes *subs*(*e*) ∈ *V*(**x**)*. For example, 〈**x**_*i,j*_, [**x**_*i,k*_, **x**_*k*+1,*j*_]〉 divides one span into two smaller ones. For each node *v*, we define its **incoming hyperedges** to be all decompositions of *v*:

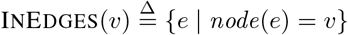

We define the **arity** of a hyperedge *e*, denoted |*e*|, to be the number of its children nodes (|*e*| ≜ |*subs*(*e*)|). In order to recursively assemble substructures to form the global structure, each hyperedge *e* = 〈*v, subs*〉 is associated with a **combination function** 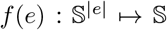 that assembles substructures from *subs* into a structure for *v* (here 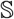 is the set of dot-bracket strings). Each hyperedge *e* is associated with an (extra) **energy term** 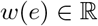.

We take the classical Nussinov algorithm (7) as a concrete example, which scores secondary structures by counting the number of base pairs. The nodes are

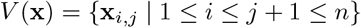

which include both non-empty substrings **x**_*i,j*_ = *x_i_*…*x_j_* (*i*≤*j*) that can be decomposed, and empty spans **x**_*i,i*−1_ (*i* = 1…*n*) that are the terminal nodes. Each non-empty span **x**_*i,j*_ can be decomposed in two ways: either base *x_j_* is unpaired (unary) or paired with some *x_k_* (*i* ≤ *k* < *j*) (binary). Therefore, the incoming hyperedges for **x**_*i,j*_ are

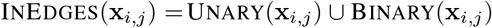

where Unary(**x**_*i,j*_) = {〈**x**_*i,j*_, [**x**_*i,j*−1_]〉} contains a single hyperedge with the combination function *f*_1_(*a*) = “*a*.” that appends an unpaired “.“ for *x_j_*:

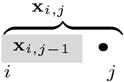

And the set of binary hyperedges

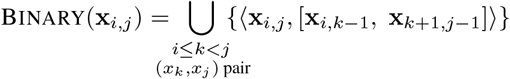

contains all bifurcations with (*x_k_, x_j_*) paired, dividing *x_i,j_* into two smaller spans **x**_*i,k*−1_ and **x**_*k*+1,*j*−1_:

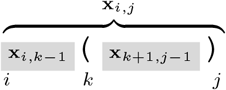

All these hyperedges share the same combination function *f*_2_(*a, b*) = “*a*(*b*)” which combines the two substructures along with the new (*x_k_, x_j_*) pair. They also share the energy term *w* = −1 (kcal/mol), the stabilizing free energy term for forming a base pair.^2^

Finally, a special **goal node** *goal*(*V*(**x**)) is identified as the root of the recursion, which in the Nussinov algorithm is the whole sequence **x**_1,*n*_.

This framework can easily extend to other folding algorithms such as Zuker (8) and LinearFold (27), where nodes are “*labeled spans*” such as *C_i,j_* for substructures over **x**_*i,j*_ with (*x_i_*, *x_j_*) paired, *M_i,j_* for multiloops over *x_i,j_*, etc.

### B. Three Sampling Algorithms

Under the hypergraph framework, we first describe the bottom-up partition function phase (also known as the “inside” or “forward” phase), and then present three algorithms for the top-down sampling phase, i.e., Non-Saving Sampling, Full-Saving Sampling, and LazySampling. While the first is similar to but cleaner than the standard sampling algorithm, the other two are novel.

#### B.0. The Partition Function Phase

In this bottom-up phase, we first calculate the local partition function *Z*(*v*) of each node *v* (see Fig. 1), summing up the contributions from each incoming hyperedge *e* (line 7) i.e., *Z*(*v*) = ∑_*e*∈InEdges_(*v*) *Z*(*e*). This part takes *O*(*E*) = *O*(*n*^3^) time as each hyperedge is traversed once and *O*(*V*) = *O*(*n*^2^) space as we need to store *Z*(*v*) for each node *v*. Note that the hyperedges are by default not saved, and will be recalculated on demand during the sampling phase. If we want to save all hyperedges (for the Full-Saving algorithm in Sec. B.2) instead, we need *O*(*n*^3^) space; the time complexity remains *O*(*n*^3^), but in practice the overhead for saving (line 8) is quite costly and it may run out of memory (see Fig. 6).

**Fig. 1.**
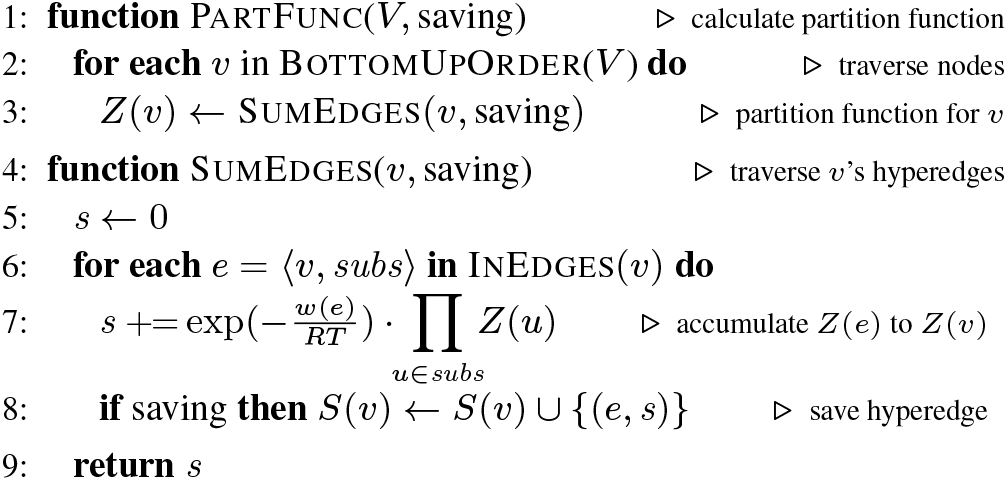
The bottom-up phase to calculate the partition function.

#### B.1. Non-Saving Sampling

In the sampling phase, Non-Saving Sampling algorithm (see Fig. 2) recursively backtraces from the goal node, in a way that is symmetric to the bottom-up partition function phase. When visiting a node *v*, it tries to sample a hyperedge *e* from *v*’s incoming hyperedges InEdges(*v*) according to the probability *Z*(*e*)/*Z*(*v*). This is done by first generating a random number *p* between 0 and *Z*(*v*), and then gradually recovering each incoming hyperedge *e*, accumulating its *Z*(*e*) to a running sum *s*, until *s* exceeds *p*, at which point that current hyperedge *e* is chosen as sampled. Note that this algorithm in general does *not* need to recover *all* incoming hyperedges of *v*, though in the worst case it would. It then backtraces recursively to the corresponding subnode(s) of hyperedge *e*.

**Fig. 2.**
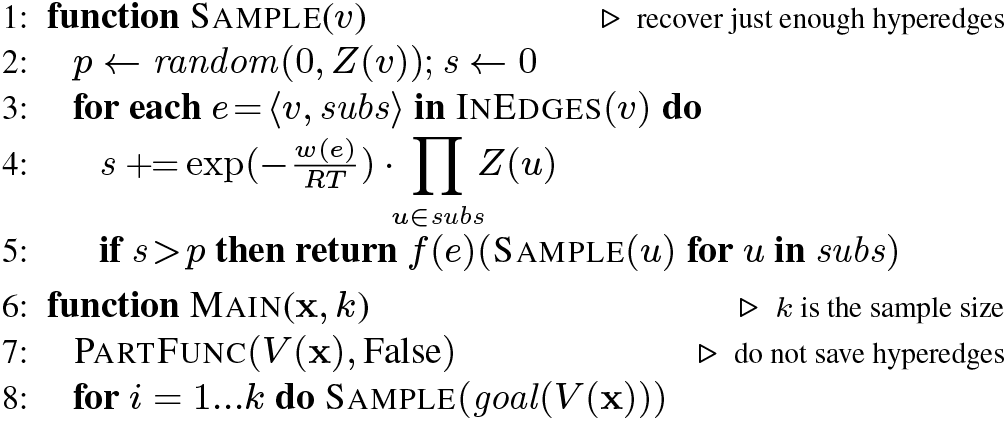
The Non-Saving Sampling algorithm. This version is similar to Ding and Lawrence (14) but simpler and cleaner thanks to the bottom-up↔top-down symmetry

Now let us analyze the time complexity to generate each sample. First of all, it visits *O*(*n*) nodes to generate one sample as there are *O*(*n*) nodes in each derivation (i.e., the recursion tree). On each node *v* = **x**_*i,j*_, it needs to recover *O*(|*j* − *i*|) hyperedges, so the total number of hypereges recovered depends on how balanced the derivation is, similar to quicksort. In the worst case (when the derivation is extremely unbalanced like a chain), it recovers *O*(*n*^2^) hyperedges, and in the best case (when the derivation is mostly balanced, i.e., bifurcations near the middle), it only recovers *O*(*n* log *n*) hyperedges. So the time to generate *k* samples is *O*(*kn*^2^) (worst-case) or *O*(*kn* log *n*) (best-case).^3^ Our experiments in Sec. 3 (Fig. SI 1) show that, like in quick sort, the sampled derivations are mostly balanced as the depth of derivation scales *O*(log *n*) in practice, thus the average case behavior is essentially best case.^4^

This version is the closest to the original Ding and Lawrence (14) algorithm, but simpler and cleaner. Our key idea is to exploit the structural symmetry between the bottom-up and sampling phases, and unify them under the general hypergraph framework. By contrast, Ding and Lawrence do *not* exploit this symmetry, and instead rely on different recurrences in the sampling phase that iteratively samples the leftmost external pair in an external span and the rightmost pair in a multiloop (see Fig. 1 of their paper). Their formulation results in unnecessarily complicated implementations (see Vienna RNAsubopt for an example).^5^ We are the first to formulate general sampling (Nussinov, Zuker, LinearFold, etc.) under a unified framework that exploits symmetry.^6^

#### B.2. Full-Saving Sampling

It is obvious that Non-Saving Sampling wastes time recovering hyperedges during the sampling phase. First, due to the symmetry, all hyperedges recovered in the sampling phase have already been traversed during the inside phase. To make things worse, many hyperedges are recovered multiple times across different samples because whenever a node is (re-)visited, its hyperedges need to be re-recovered. This situation worsens with the sample size *k*. More formally, we define

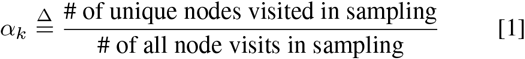

to be the “unique visit ratio” among *k* samples, and we will see in Fig. 8A that this ratio is extremely small, quickly approaching 0% as *k* increases, meaning most node visits are repeated visits. This begs the question: why don’t we save all hyperedges during the inside phase, so that no hyperedge needs to be recovered during the sampling phase? To address this we present the second, Full-Saving Sampling (see Figs. 2–3), which saves for each node *v* the contributions *Z*(*e*) of each hyperedge *e* to the local partition function *Z*(*v*), once and for all. Then the sampling phase is easier, only requiring sampling a hyperedge *e* according to its relative contribution (or “weight”) to *v*, i.e., *Z*(*e*)/*Z*(*v*) (line 2 in Fig. 3). Actually, modern programming languages such as C++ and Python provide tools for sampling from a weighted list, which is implemented via a binary search in the sorted array of cumulative weights (which is why line 8 in Fig. 1 saves the running sum rather than individual contribution *Z*(*e*)). This takes only *O*(log *n*) time for each *v* as |InEdges(*v*)| = *O*(*n*) (consider all bifurcations). Therefore, the worst-case complexity for generating *k* samples is *O*(*kn* log *n*) and the best-case is *O*(*kn*).^7^

**Fig. 3.**
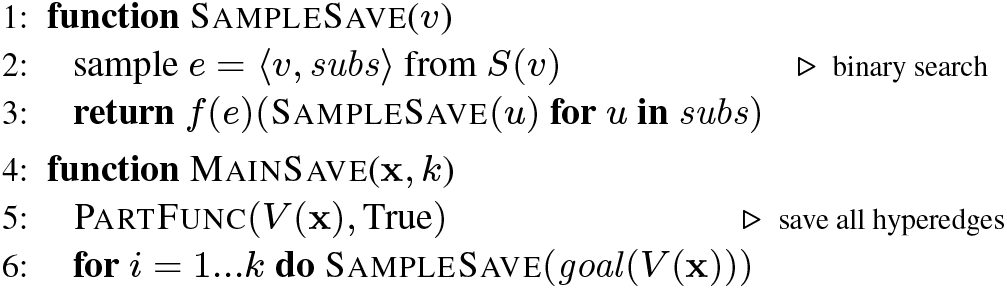
The Full-Saving Sampling algorithm.

#### B.3. LazySampling = Lazy-Saving Sampling

Though Full-Saving Sampling avoids all re-calculations, it costs too much more space (*O*(*n*^3^) vs. *O*(*n*^2^)) and significantly more time in practice for saving the whole hypergraph. Actually, the vast majority of nodes are *never* visited during the sampling phase even for large sample size. To quantify this, we define

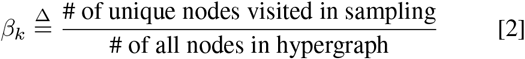

to be the “visited ratio”. Our experiments in Fig. 8B show that only < 0. 5% of all nodes in the hypergraph are ever visited for 20,000 samples of a 3,048 *nt* sequence using Vienna RNAsubopt, so most of the saving is indeed wasted. Based on this, we present our third version, LazySampling, which is a hybrid between Non-Saving and Full-Saving Samplings (see Fig. 4). By “lazy” we mean only recovering and saving a node *v*’s hyperedges when needed, i.e., the first time *v* is visited during sampling phase. In this way each hyperedge is recovered at most once, and most are not recovered at all. This version balances between space and time, and is the fastest among the three versions in most settings in practice.

**Fig. 4.**
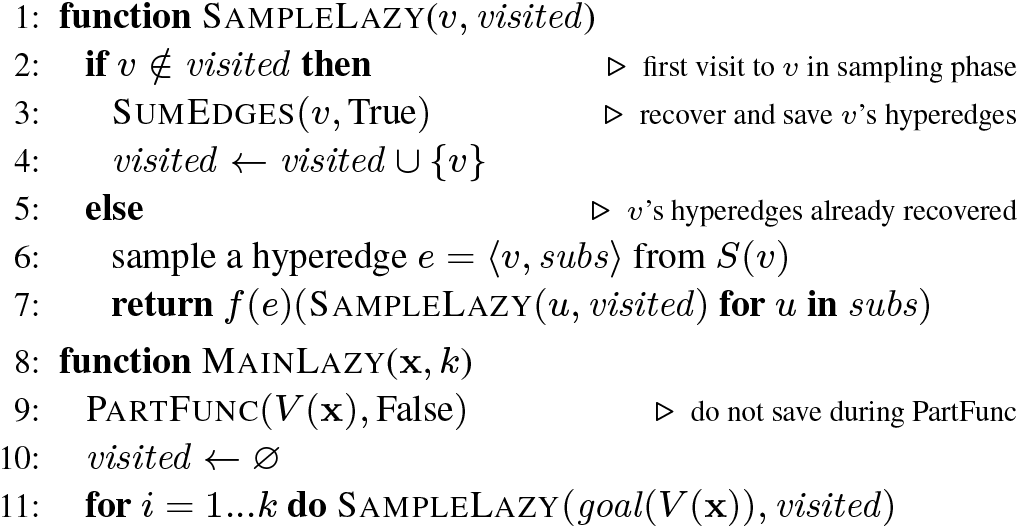
The LazySampling (Lazy-Saving Sampling) algorithm.

The complexity analysis of LazySampling is also a hybrid between the Non- and Full-Saving versions, combined together using the *α_k_* and *β_k_* ratios. We note that the sampling time of LazySampling consists of two parts: (a) the hyperedge-recovering (and saving) work, and (b) the sampling work after relevant hyperedges are recovered. Part (a) resembles Non-Saving Sampling, but with a ratio of *α_k_*, because most node visits are repeated ones, and once a node is visited for the first time in sampling, its hyperedges are recovered and saved, and all future visits to this node will be like full saving version. Part (b) is identical to Full-Saving Sampling (in both cases, all needed hyperedges are already saved). Therefore, we have the following relations among the time complexities for the sampling phase of these three versions:

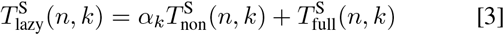

This holds for both the worst and best-case scenarios in Tab. 1. The space complexity is easier: LazySampling saves only a fraction (*β_k_*) of all nodes in the hypergraph, thus *O*(*β_k_n*^3^). See Tab. 1 for summary.

**Table 1.** Complexities for four sampling algorithms (*n* is the sequence length, *k* is sample size, *α_k_* and *β_k_* are the “unique visit ratio” (Eq. 1) and “visited ratio” (Eq. 2), resp., and *b* is beam size). The runtimes of LazySampling (i.e., Lazy-Saving Sampling) are hybrids between the Non- and Full-Saving ones, i.e.,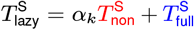 (Eq. 3). LinearSampling extends LazySampling by replacing the exact partition function with LinearPartition, and achieves end-to-end linear runtime and space against *n*.

| sampling algorithm | partition function time | function space | sample time (worst-case) | sample time (best-case) | sample space (worst-case) |
| --- | --- | --- | --- | --- | --- |
| Non-Saving Sampling | $O(n^3)$ | $O(n^2)$ | $O(kn^2)$ | $O(kn \log n)$ | $O(1)$ |
| Full-Saving Sampling | | $O(n^3)$ | $O(kn \log n)$ | $O(kn)$ | |
| LazySampling | | $O(n^2)$ | $O(\alpha_k kn^2 + kn \log n)$ | $O(\alpha_k kn \log n + kn)$ | $O(\beta_k n^3)$ |
| LinearSampling | $O(nb^2)$ | $O(nb)$ | $O(\alpha_k knb + kn \log b)$ | $O(\alpha_k kn \log b + kn)$ | $O(\beta_k nb^2)$ |

### C. LinearSampling = LazySampling + LinearPartition

LazySampling is the most efficient among all three methods presented above, but the biggest bottleneck remains the *O*(*n*^3^)-time partition function computation, which prevents it from scaling to full-length viral genomes such as SARS-CoV-2. To address this problem, we replace our recently proposed linear-time approximate algorithm, LinearPartition (23), to replace the standard cubic-time one. It can be followed by any one of the three sampling algorithms (Non-Saving, Full-Saving, and LazySampling) for the sampling phase, and in particular, we name the one with Lazy-Saving the LinearSampling algorithm as it is the fastest among all combinations.

Fig. 5 describes a simplified pseudocode using the Nussinov-Jacobson energy model. Inspired by LinearPartition, we employ beam search to prune out nodes with small partition function (line 11) during the inside phase. So at each position *j*, only the top *b* promising nodes “survive” (i.e., *O*(*nb*) nodes survive in total). Here the beam size *b* is a user-specified hyperparameter, and the default *b* = 100 is found to be more accurate for structure prediction than exact search (23). The partition function runtime is reduced to *O*(*nb*^2^) (there are only *O*(*b*) hyperedges per node) and the space complexity is reduced to *O*(*nb*), both of which are linear against sequence length *n*. The sampling time is also linear, and the binary search time to sample a saved hyperedge reduces from *O*(log *n*) to *O*(log *b*) since at most *b* hyperedges are saved for each node (thanks to beam search). Then, following Eq. 3, we can derive the complexities in Tab. 1. In particular, the LinearSampling algorithm (the last line in the table) has an end-to-end runtime of *O*(*nb*^2^ + *α_k_kn* log *b* + *kn*) and uses *O*(*nb* + *β_k_nb*^2^) space in total, both of which scales linearly in *n*.

**Fig. 5.**
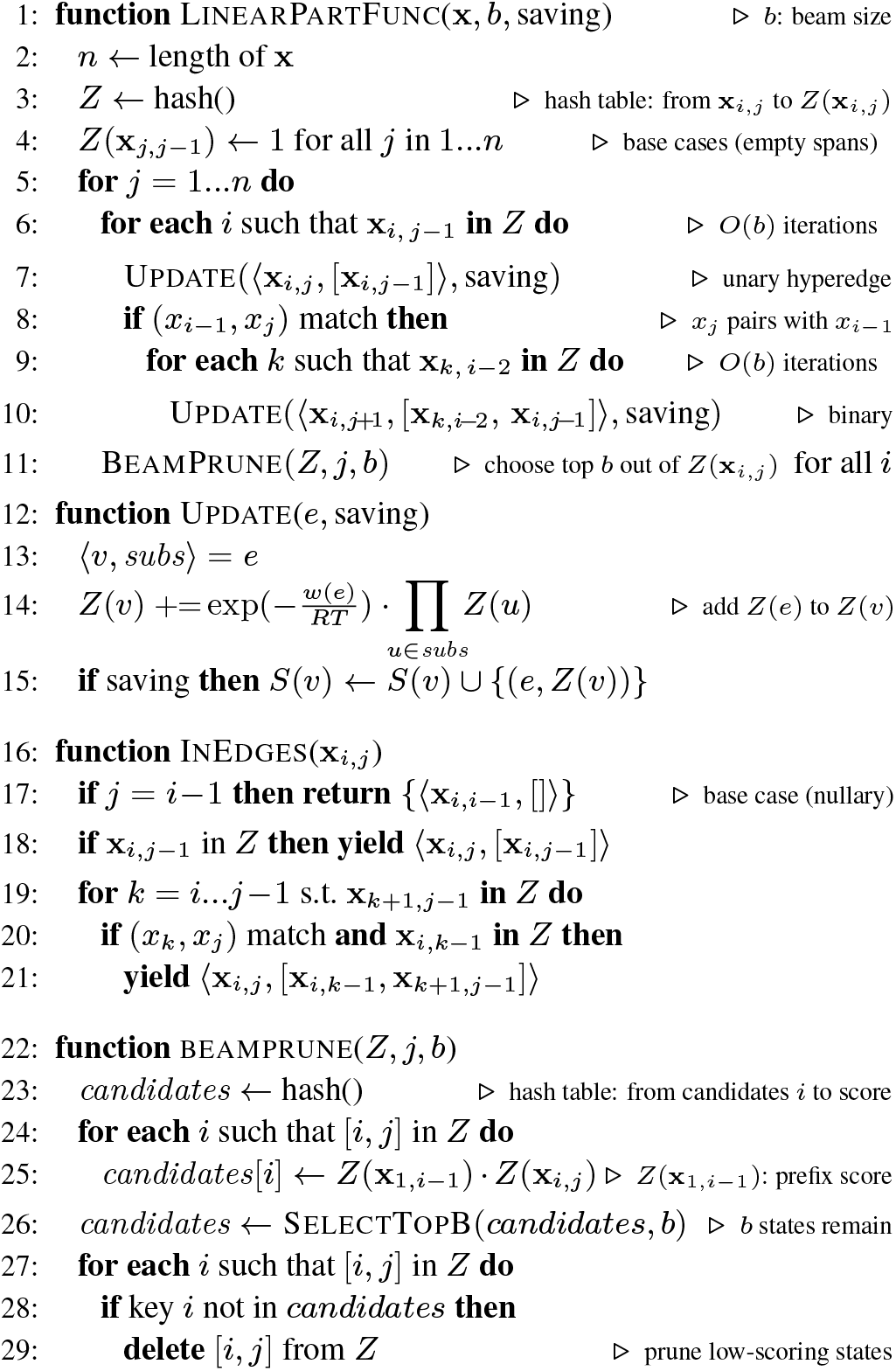
Part of the (simplified) pseudocode of linear-time sampling algorithms on the Nussinov-Jacobson energy model, which can be called by the Sample (Fig. 2), SampleSave (Fig. 3), and SampleLazy (Fig. 4) functions, by replacing PartFunc in those functions with LinearPartFunc here. The InEdges function is used in the SumEdges function in the non-saving and lazy-saving versions. The actual algorithm using the Turner model is available on https://github.com/LinearFold/LinearSampling.

## 3. Results

### A. Efficiency and Scalability

We benchmark the runtime and memory usage on 26 sequences sampled from RNAcentral (32). We evenly split the range from 0 to 8,000 into 52 bins by log-scale, and randomly select at most one sequence in each bin; within 100 *nt* only one sequence is chosen. We refer this dataset as the RNAcentral dataset in the paper. We use a Linux machine (CentOS 7.5.1804) with 3.40 GHz Intel Xeon E3-1231 v3 CPU and 16 GB memory, and gcc 4.8.5.

#### A.1. Comparing Non-Saving, Full-Saving, LazySampling and LinearSampling

Fig. 6 shows the performance of different versions of sampling algorithms, using Vienna RNAsubopt (19) as a baseline. Note that RNAsubopt, Non-Saving, Full-Saving and LazySampling are under the exact partition function calculation, while LinearSampling uses linear partition function. Our own exact partition function (basically, setting *b* = +∞) is faster than RNAsubopt with identical results. Regarding end-to-end runtime, Full-Saving Sampling is the slowest since it spends much time on hyperedges saving, and it runs out of memory on a 3,048 *nt* sequence. Non-Saving Sampling is much faster than Full-Saving, but slightly slower than LazySampling; both Non-Saving and LazySampling are faster than RNAsubopt. The partition function runtime is close to end-to-end. For sampling-only time, LazySampling is similar to Full-Saving Sampling, and is more than 2.5× faster than Non-Saving and RNAsubopt. Regarding memory usage, Full-Saving uses much more memory, while the other three are close. Saliently, by integrating a linear partition function to LazySampling, LinearSampling significantly reduces the partition function time and memory usage, and is also the fastest in sampling phase.

**Fig. 6.**
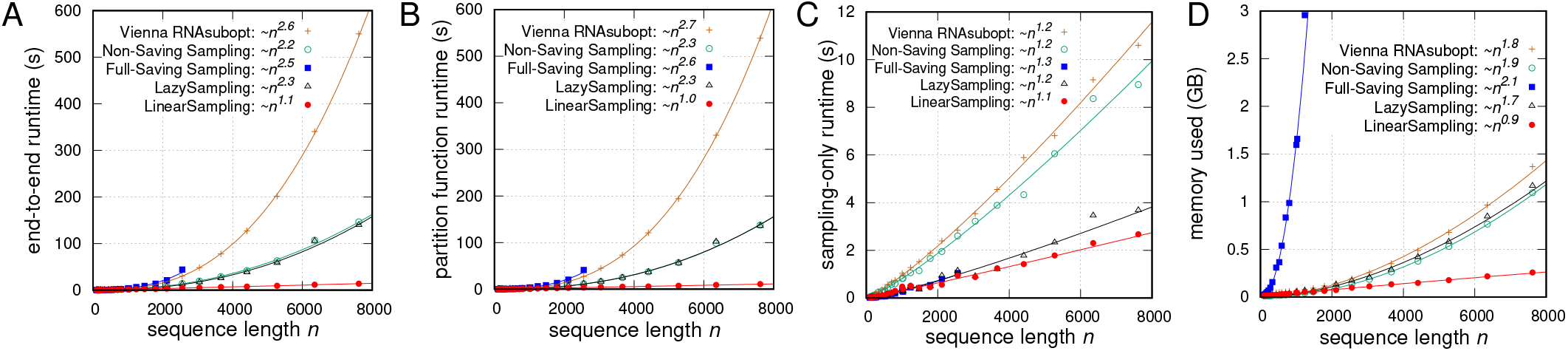
Runtime and memory comparisons against sequence length *n* on RNAcentral dataset, using exact partition function (*k* = 10,000 samples). **A**: the end-to-end runtime comparison. **B**: the partition function (inside phase) runtime comparison. **C**: the sampling-only (outside phase) runtime comparison. **D**: the memory usage comarison; the y-axis is cut at 3 GB to show the data points more clear. Note that Full-Saving Sampling runs out of memory on the 3,048 *nt* sequence. The runtime and memory comparisons of different sampling strategies with linear partition function are shown in Fig. SI 2.

The performance of the sampling algorithms also depend on the sample size *k*, so in Fig. 7, we present the comparisons against *k* on a 2,558 *nt* sequence, which is the longest one that Full-Saving Sampling can finish with the memory limit in the dataset. End-to-end, Full-Saving Sampling is about 3× slower than Non-Saving with a small *k*, but with the increase of *k* the gap shrinks. LazySampling has identical runtime to Non-Saving in partition function phase and similar runtime to Full-Saving in sampling phase, and has the smallest end-to-end runtime among these three versions. The subfigure in Fig. 7C zooms in the sampling-only time in small sample size, where LazySampling is slightly slower than Non-Saving when *k* < 340, but is faster otherwise. It is consistent with our observation that the cost of recovering and storing hyperedges on demand in LazySampling is small, and suggests to use LazySampling in stead of Non-Saving. RNAsubopt uses the non-saving strategy, so its performance trend is similar to our Non-Saving Sampling, but much slower; while LinearSampling uses the lazy-saving strategy and it is similar to LazySampling but much faster.

**Fig. 7.**
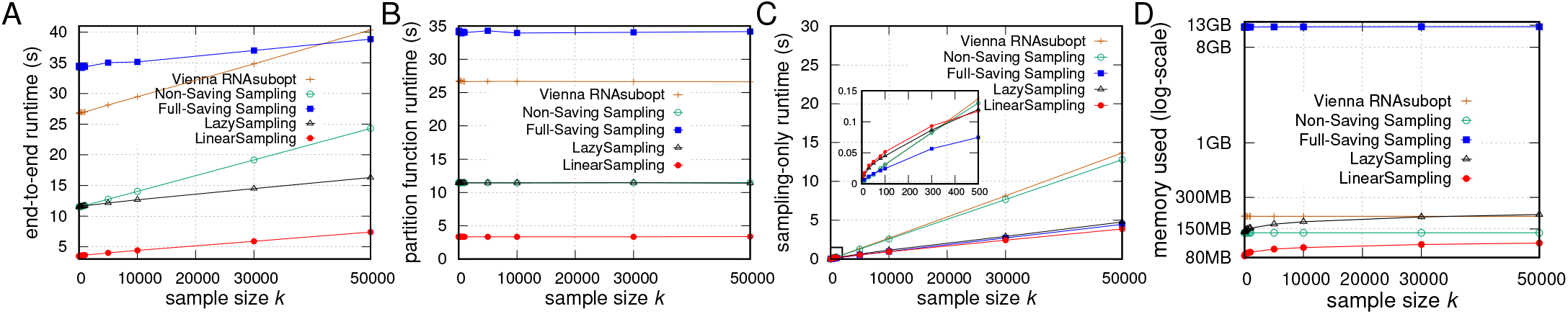
Runtime and memory usage comparisons against sample size *k* on a 2,558 *nt* sequence. **A**: the end-to-end runtime comparison. **B**: the partition function (inside phase) runtime comparison. **C**: the sampling-only (outside phase) runtime comparison; here we zoom in 0 ≤ *k* ≤ 500 interval. **D**: the memory usage comarison; the y-axis is in log-scale. The runtime and memory comparisons of different sampling strategies with linear partition function against *k* are shown in Fig. SI 3.

We also illustrate why LazySampling is a better sampling strategy in practice. Fig. 8A shows the unique visit ratio *α_k_* is less than 5% when *k* > 1,000 for both RNAsubopt and LinearSampling, confirming that Lazy-Saving is able to avoid a large number of re-calculations during the sampling phase. On the other hand, the visited ratio *β_k_* (Fig. 8B) is always smaller than 0.5% and 3% for RNAsubopt and LinearSampling, resp, and grows slower and slower as the sampling size is increasing, showing that saving all hyperedges (i.e., Full-Saving) is not ideal. Fig. 8C and D further confirm that both *α_k_* and *β_k_* do not increase with sequence length, therefore this analysis applies to both short and long sequences.

**Fig. 8.**
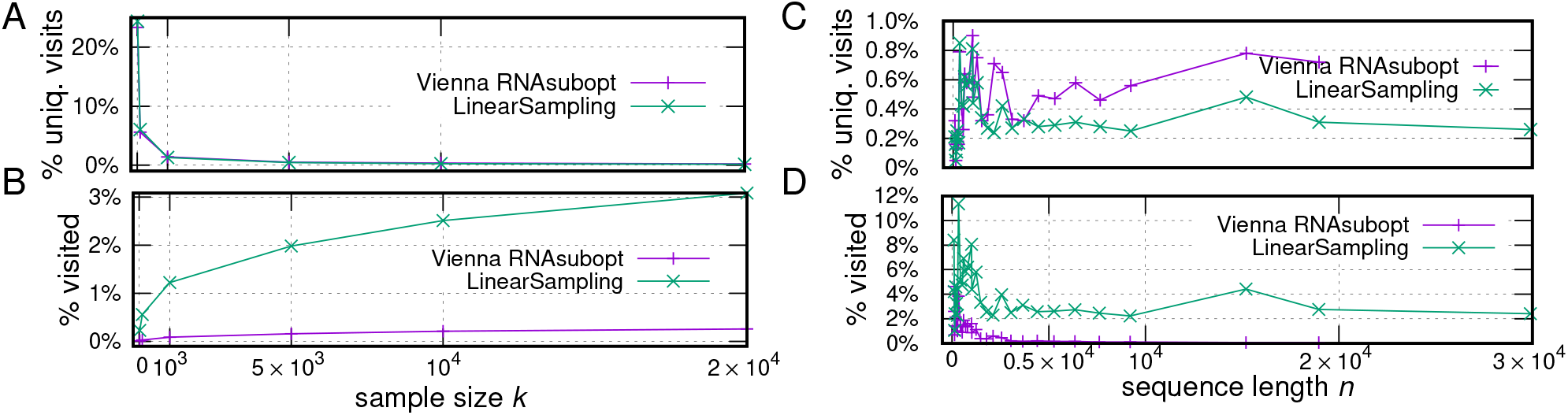
In practice most node visits are repeated, and only a small portion of nodes are visited. The sample size is 1,000. **A** and **B**: unique visit ratio *α_k_* and visited ratio *β_k_* against sample size *k*; here *n*=3,048 *nt*. **C** and **D**: *α_k_* and *β_k_* against sequence length *n*; here *k*=10,000. Fig. SI 4 demonstrates that most of the visits are concentrated on a few nodes.

#### A.2. Comparing LinearSampling to Vienna RNAsubopt Global and Local Modes

We compare the efficiency and scalability between LinearSampling and RNAsubopt. To investigate their performance on long sequences, e.g., full-length viral genomes, we extend our benchmark dataset by adding two longer sequences (19,071 *nt* and 22,158 *nt*) from RNAcentral, as well as four viral sequences, HIV (9,181 *nt*), RSV (15,191 *nt*), Ebola (18,959 *nt*) and SARS-CoV-2 (29,903 *nt*).

Regarding end-to-end runtime, LinearSampling scales almost linearly against sequence length, and is much faster than RNAsubopt. LinearSampling is 176× faster (38.9s vs. 1.9h) than RNAsubopt on the full genome of Ebola virus (18,959 *nt*), and can finish the full-length of SARS-CoV-2 in 69.2s; while RNAsubopt runs out of memory on SARS-CoV-2. Regarding sampling-only runtime, LinearSampling is more than 3× faster. Fig. 9C confirms that the memory usage of LinearSampling is linear, but RNAsubopt requires *O*(*n*^2^) memory. LinearSampling uses less than 1 GB memory for Ebola sequence, while RNAsubopt uses more than 8 GB.

**Fig. 9.**
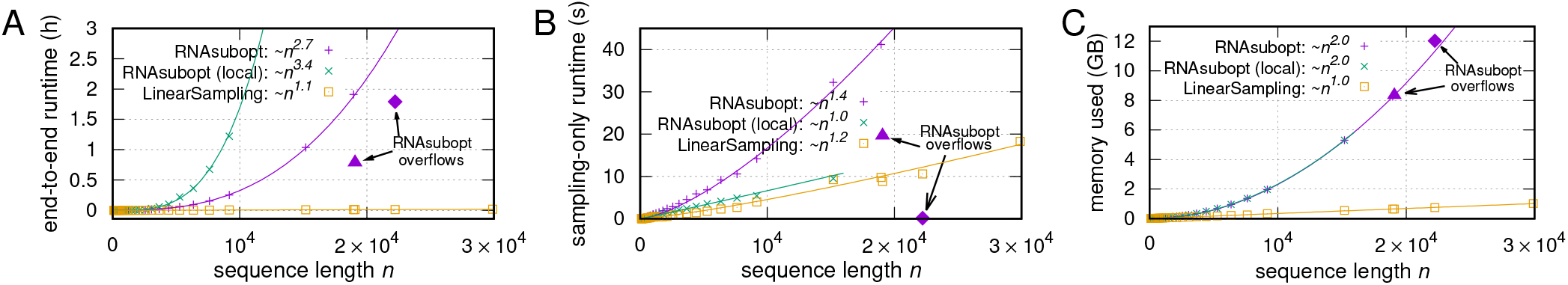
Runtime and memory usage comparisons between RNAsubopt, its local mode (span=70), and LinearSampling on the RNAcentral dataset and viral sequences. The sample size is 10,000. LinearSampling can scale to the full genome of SARS-CoV-2 (~30,000 *nt*), but both RNAsubopt global and local modes cannot. The purple triangles and diamonds are the sequences that RNAsubopt overflows.

It is surprising that RNAsubopt local mode has a complexity of *O*(*n*^3,4^) for end-to-end runtime, and is even slower that its global mode. For a 9,181 *nt* sequence, RNAsubopt local mode takes 73.2 minutes, and its global mode takes 15.2 minutes. As a comparison, LinearSampling only takes 18 seconds. Memory-wise, RNAsubopt local mode uses as much as its global mode. The benchmark experiment suggests that RNA-subopt local mode is not able to scale up to long sequences.

We also observe that, similar to RNAfold, RNAsubopt sometimes overflows on long sequences during the partition function calculation (23), making it less reliable for long sequences. For example, RNAsubopt overflows on two sequences from RNAcentral, shown in Fig. 9 with purple triangles and diamonds. For the triangle one (URS00007C400D, 19,071 *nt*), an overflow happens in the segment [5775, 12619], leading to an unpaired region longer than 6,000 *nt* in all sampled structures. It is clear in Fig. 9 that both the end-to-end and sampling-only runtimes drop unreasonably. For the diamond one (URS00009C28A8_9606, 22,158 *nt*), the overflow triggers an error during sampling phase, “ERROR: backtrack failed in qm1”, resulting in an abnormal exit of the software, with only a few structures generated. We can see that though self-adapted scaler is used in RNAsubopt, overflow is unavoidable for some long sequences. As a contrast, LinearSampling uses log-space for partition function, and does not have overflow issue.

### B. Quality of the Samples

We use the ArchiveII dataset (29; 33), which contains a set of sequences with well-determined structures, to investigate the quality of the samples. We follow the preprocessing steps of a previous study (23), and obtain a subset of 2,859 sequences distributed in 9 families.

#### B.1. Approximation Quality to Base Pairing Probabilities

To evaluate if the sampling structures approximate to the ensemble distribution, Ding and Lawrence (14) investigated the frequency of the MFE structure appeared in the samples, and checked if it matches with the Boltzmann distribution. However, this only works for short sequences because the frequency of the MFE structure is extremely small for long sequences, e.g., 2.23×10^−32^ for *E. coli* 23S rRNA (around 3,000 *nt*). Alternatively, we investigate the root-mean-square deviation (**RMSD**) between the base pairing probability matrices *p*(*S*), which is derived from the sample set *S*, and *p′*, which is generated by Vienna RNAfold or LinearPartition. Note that rmsd is averaged on all possible Watson-Crick and G-U pairs on the sequence (34).

Fig. 10A shows four curves of average rmsd of base pairing probabilities against sample size on the ArchiveII dataset. The green curve illustrates the rmsd between LinearSampling and RNAfold. Although LinearSampling approximates the partition function based on LinearPartition, which introduces small changes in base pairing probabilities (23), the rmsd is only 0.015 with *k* = 10, and drops down to 0.005 quickly with sample size 5,000. As a comparison, RNAsubopt local mode (with a base pair distance limit of 150) has a larger rmsd when approximating to RNAfold, shown in the red curve. Regarding the rmsd between LinearSampling and LinearPartition (the blue curve), and between RNAsubopt and RNAfold (the yellow curve), we observe that they are almost identical, suggesting LinearSampling can generate structures strictly matching with the ensemble distribution as well as RNAsubopt.

**Fig. 10.**
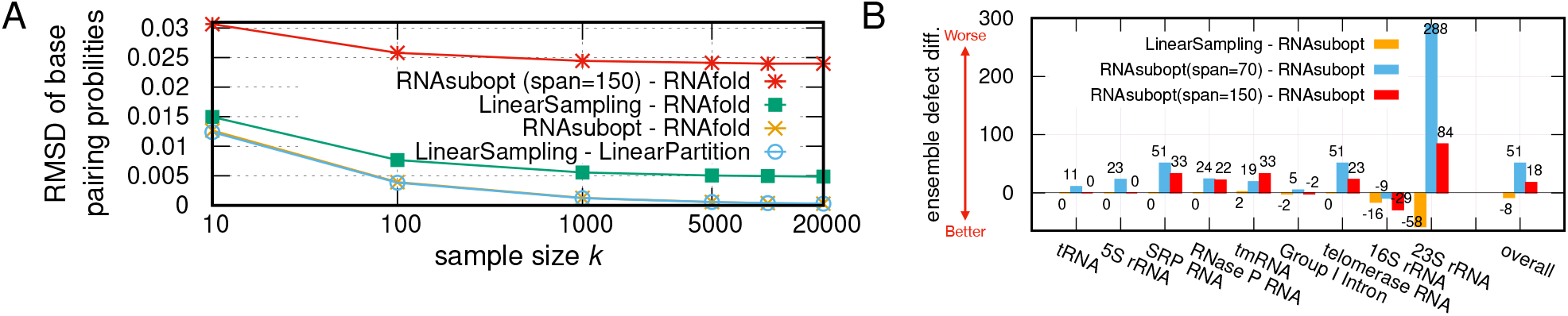
LinearSampling matches with the ensemble distribution, and is better correlated with the ground truth structure. **A**: the root-mean-square deviation (RMSD) of base pairing probabilities against sample size. The RMSD is averaged within each family, and then averaged on all families. **B**: the ensemble defect difference of each family and overall (averaged by families), between LinearSampling and RNAsubopt, and between RNAsubopt local modes (span=70 and 150, respectively) and RNAsubopt global mode.

#### B.2. Correlation with the Ground Truth Structure

We investigate the sampled structure’s correlation with the ground truth using “ensemble defect” (35), the expected number of incorrectly predicted nucleotides over the ensemble. It is defined:

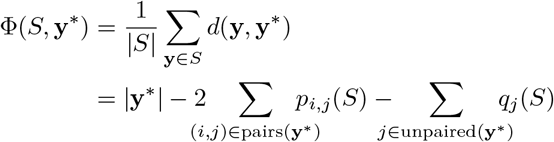

where **y**\* is the ground truth structure, and *d*(**y**, **y**\*) is the distance between **y** and **y**\*, defined as the number of incorrectly predicted nucleotides in **y**. And *q_j_*(*S*) is the probability of *j* being unpaired in the sample *S*, i.e., *q_j_*(*S*) = ∑*p_i,j_*(*S*).

Fig. 10B shows the ensemble defect differences between LinearSampling and RNAsubopt (orange bars) on each family (ordered in their average sequence length, from the shortest to the longest) and overall. Note that better correlation to the ground truth structures requires lower ensemble defect. For short families, the differences between LinearSampling and RNAsubopt are either 0 or close to 0, indicating that the sampling qualities of the two systems are similar on these families. But on 16S and 23S rRNAs, LinearSampling has lower ensemble defect, showing that it performs better on longer sequences. The only family that LinearSampling performs worse is tm-RNA. We also present the comparisons between RNAsubopt local and global modes, with base pair length limitations of 70 (blue bars) and 150 (red bars).^8^ It is obvious that the local sampling has much higher (worse) ensemble defect on 23S rRNA, which may due to the ignore of all base pairs beyond the max span limit.

An important application of the sampling algorithm is to calculate a region’s accessibility.^9^ Therefore, we calculate accessibilities of window size 4 (14) from structures generated by LinearSampling and RNAsubopt, as well as directly from RNAplfold (38), and evaluate based on the ground truth structures. We denote the measurement of *accessibility defect* as *D*, which evaluates the averaged wrong predictions of the accessibility to the ground truth given a window size. For sampling-based methods, *D*(*S*, **y**\*) is generated from the samples *S* and is defined as:

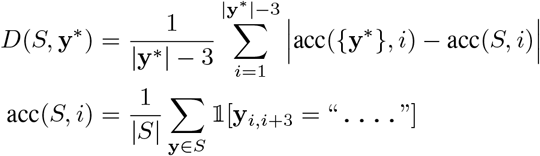

where acc(*S, i*) is the accessibility of region [*i, i* + 3].

Table 2 shows the accessibility defect comparison on ArchiveII. We observe that LinearSampling outperforms (or is as good as) all the other systems on 7 out of 9 families, and is the best overall. The only two families which LinearSampling is not the best are tmRNA and Group I Intron, for which LinearSampling is still among the top three. Notably, both RNAsubopt’s and RNAplfold’s local modes are normally worse than their global modes, with only one exemption on Group I Intron family, indicating that the local modes are less accurate.

**Table 2.** Accessibility defect comparison between Vienna RNAsubopt, the RNAsubopt local sampling and LinearSampling on the ArchiveII dataset. The sample size is 10,000. The best (lowest) accessibility in each collumn is highlighted in red.

|  | tRNA | 5S rRNA | SRP RNA | RNase P RNA | tmRNA | Group I Intron | telomerase RNA | 16S rRNA | 23S rRNA | overall |
| --- | --- | --- | --- | --- | --- | --- | --- | --- | --- | --- |
| LinearSampling | **0.1823** | **0.1552** | **0.1309** | **0.2125** | 0.2249 | 0.2637 | **0.2994** | **0.1804** | **0.1371** | **0.1985** |
| RNAsubopt | 0.1826 | 0.1553 | **0.1309** | **0.2125** | **0.2245** | 0.2649 | 0.2998 | 0.1841 | 0.1405 | 0.1995 |
| RNAsubopt (span=70) | 0.2237 | 0.2097 | 0.1925 | 0.2380 | 0.2437 | 0.2623 | 0.3259 | 0.2006 | 0.1710 | 0.2297 |
| RNAsubopt (span=150) | 0.1825 | 0.1553 | 0.1475 | 0.2270 | 0.2423 | **0.2607** | 0.3091 | 0.1828 | 0.1528 | 0.2067 |
| RNAplfold | 0.1825 | 0.1553 | **0.1309** | **0.2125** | **0.2245** | 0.2649 | 0.2998 | 0.1993 | 0.1617 | 0.2035 |
| RNAplfold (span=70) | 0.2249 | 0.2348 | 0.2132 | 0.2638 | 0.2693 | 0.2986 | 0.3255 | 0.2355 | 0.2103 | 0.2529 |
| RNAplfold (span=150) | 0.1825 | 0.1553 | 0.1561 | 0.2426 | 0.2520 | 0.2761 | 0.3207 | 0.2060 | 0.1771 | 0.2187 |

It is worth noting that the better results of ensemble defect and accessibility defect of LinearSampling are inherited from LinearPartition, which correlates better with the ground truth structures (23).

### C. Applications to SARS-CoV-2

The COVID-19 pandemic has swept the world in 2020 and 2021, and is likely to be a threaten of global health for a long time. Therefore, it is of great value to find the regions with high accessibilities in SARS-CoV-2, which can be potentially used for COVID-19 diagnostics and drug design. But since SARS-CoV-2 is as long as 30,000 *nt*, all existing computational tools are unable to be applied to its full-length genome globally. Now with significant improvement on sampling efficiency and scalability,LinearSampling is able to fast sample structures for the wholegenome of SARS-CoV-2, and predict its accessible regions.

We run LinearSampling on NC_0405512.2, the reference sequence of SARS-CoV-2 (39). We also take RNAplfold local mode (span=150) as a baseline. RNAsubopt local mode is too slow and out of memory on SARS-CoV-2, so we does not include it in the experiment. First, we check if the accessibilities predicted by LinearSampling match better with the experimentally-guided structures, especially on the regions of interest, e.g., the 5’-UTR region which has conserved structures and plays a critical role in viral genome replication (40). Fig. 11 compares the accessibilities derived from LinearSampling and RNAplfold to the experimentally-guided structures based on SHAPE reactivities (24). Following Ding and Lawrence (14), the accessibilities of window size 4 are visualized in the top sub-figure, and LinearSampling clearly correlates better with the SHAPE-directed structure. For example, RNAplfold overestimates the accessibilities around the double-standed region [25, 33], while LinearSampling’s predictions are close to 0. Also, LinearSampling correctly captures the accessible region around 50, with a high predicted accessibility of nearly 1. We further extend the window size to cover a wider range (from 1 to 11), and visualize the results in the middle (LinearSampling vs. SHAPE-directed) and bottom (RNAplfold vs. SHAPE-directed) sub-figures. For instance, the red circle at position 66 and window size 7 (pointed by a red arrow), representing a highly accessible region [66, 72] predicted by LinearSampling, is surrounded by a box, which indicates that the prediction is supported by the wetlab experiment. In RNAplfold’s prediction, the same position (pointed by a blue arrow) is in yellow, indicating that it has lower accessibility and is less correlated with the SHAPE reactivities. The main differences between LinearSampling and RNAplfold are highlighted in gray shades. In general, LinearSampling’s result correlates better with the experimentally-guided structure.

**Fig. 11.**
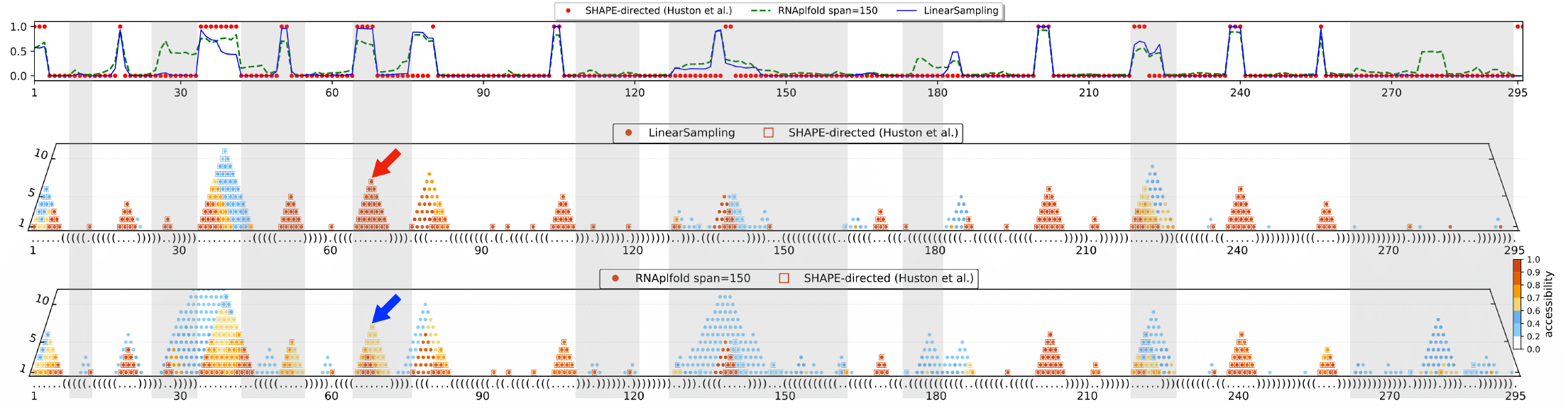
The accessibilities derived from LinearSampling correlate better with the unpaired region in the SHAPE-directed structure of SARS-CoV-2 5’-UTR (24). Note that the full sequence was used for the accessibility calculation, but we only illustrate the 5’-UTR region as an example. **Top**: accessibilities of window size 4 derived from SHAPE-directed structure (red dots), RNAplfold local mode (dashed green line) and LinearSampling (solid blue line). Following Ding and Lawrence (14), the accessibility of position *i* stands for the region [*i, i* + 3]. **Middle**: accessibilities predicted by LinearSampling with window sizes from 1 to 11. Each prediction is presented with a solid circle, where the color correlates with its accessibility value. The SHAPE-directed structure from Huston et al. (24) is shown in dot-bracket format along the x-axis, and its accessible regions are annotated in boxes. **Bottom**: accessibilities predicted by RNAplfold local mode with a base pairing limit of 150. Note that the top sub-figure is a special case (window size 4) of the bottom two sub-figures. The main differences between the predictions of LinearSampling and RNAplfold are highlighted in gray shades. The red and blue arrows point to position 66 and window size 7, which represents the region [66, 72].

To further quantify the difference between LinearSampling and RNAplfold, we calculate the accessibility defects of three important regions in SARS-CoV-2, 5’-UTR, the Frameshifting Element (FSE), and 3’-UTR, shown in Table 3. LinearSampling has lower (better) accessibility defects on all these three regions, suggesting that LinearSampling is a more reliable tool for SARS-CoV-2 study.

**Table 3.** Accessibility defect comparisons between LinearSampling and RNAplfold on UTRs and Frameshifting Element (FSE).

|  | 5'-UTR | FSE | 3'-UTR |
| --- | --- | --- | --- |
| LinearSampling | **0.0757** | **0.0926** | **0.2203** |
| RNAplfold (span=150) | 0.1324 | 0.1502 | 0.2941 |

Secondly, we aim to obtain potentially accessible regions. A previous study (41) locates conserved unstructured regions of SARS-CoV-2 by scanning the reference sequence with windows of 120 *nt*, sliding by 40 *nt*, and then calculating base pairing probabilities using CONTRAfold (42) for these fragments. In total, 75 accessible regions with 15 or more nucleotides are claimed, where each base has the average unpaired probability of at least 0.6. However, this method has two flaws: (1) it is *not* correct principally to evaluate accessibility based on unpaired probabilities due to their mutual dependency; and (2) it neglects long-range base pairs and has to approximate the accessibilities based on local structures.

Instead, we measure the accessibilities based on samples generated by LinearSampling, setting the window size to be 15 following Rangan et al. (41). We only show the fragments whose accessibilities are larger than 0.5, i.e., they are more likely to be opening than closing. We list all 23 regions found by LinearSampling in Table 4. Some of the regions are overlapped, resulting in a total of 9 separate accessible regions, which are illustrated in Fig. 12. Among the 9 regions, two are in ORF1ab, one in ORF3a, one in the M gene, three in the N gene, and two in the S (spike) gene, whose proteins can recognize and bind with receptor (43).

**Table 4.**
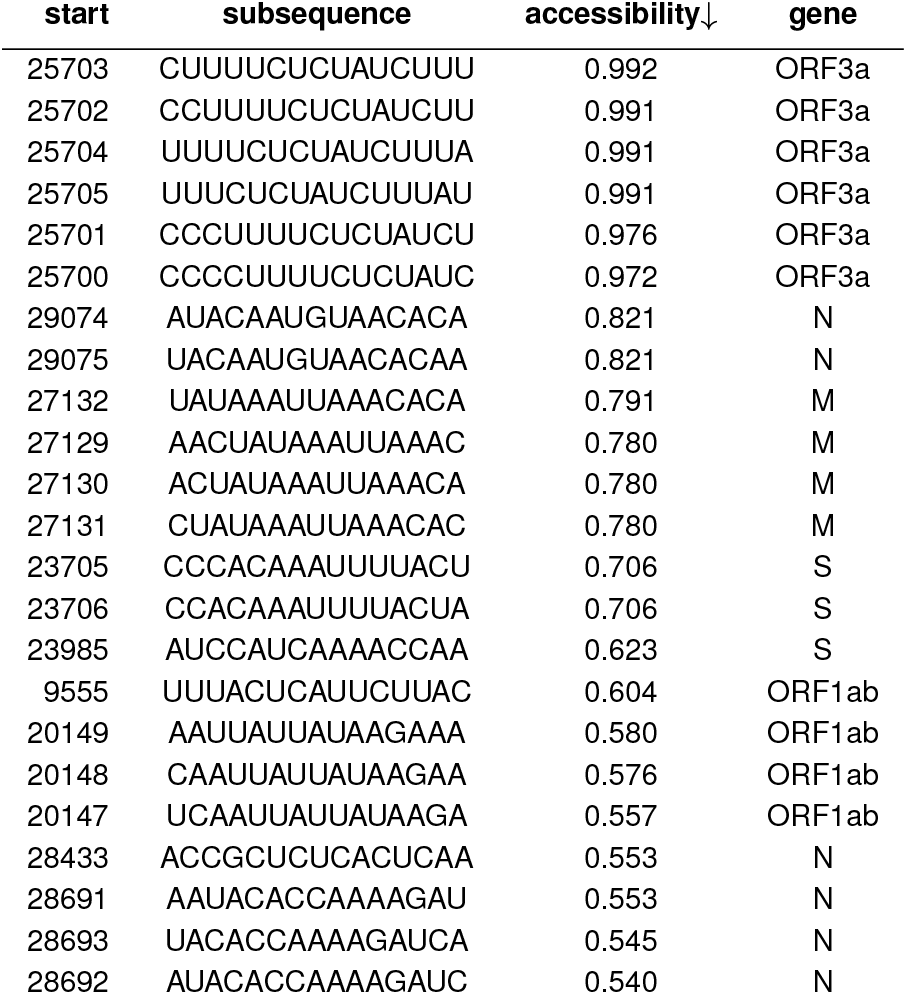
Regions of 15 nucleotides with negative log-odds of free energy change. They are found by LinearSampling running on the SARS-CoV-2 full genomes.

**Fig. 12.**
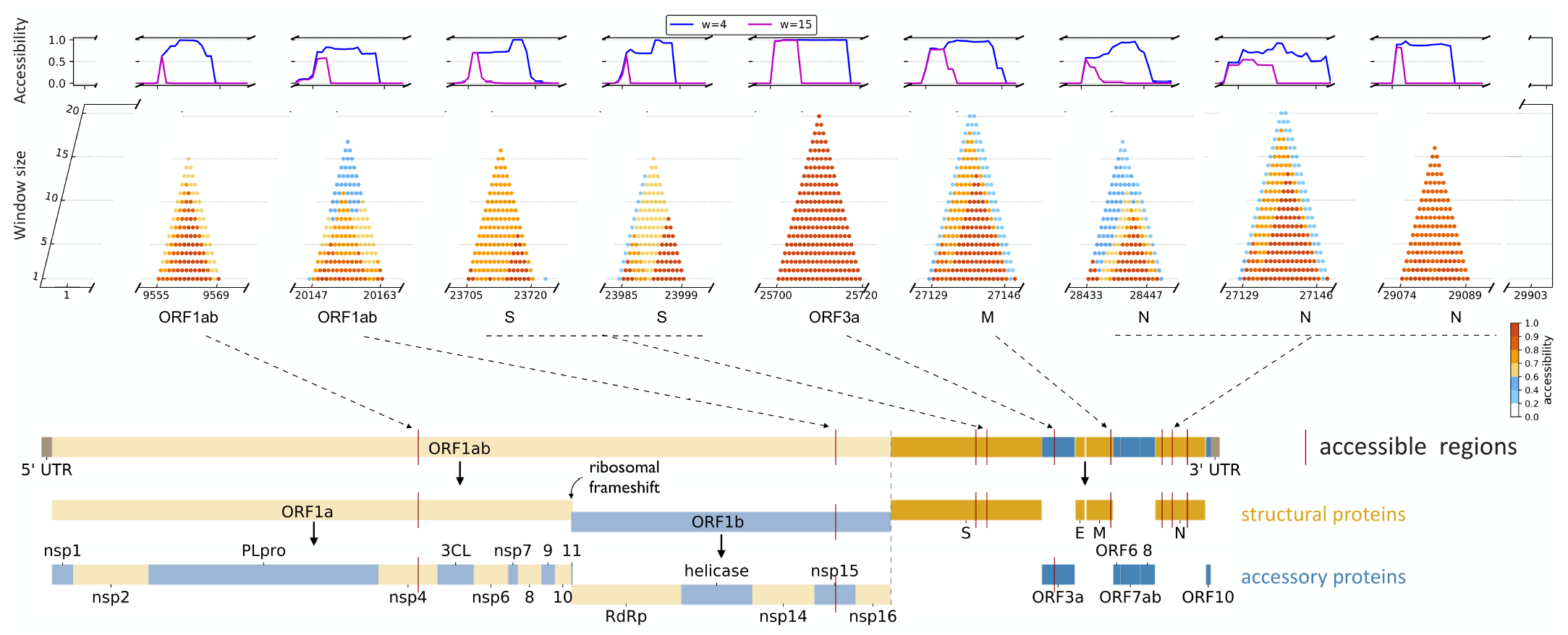
LinearSampling predicts 9 separate accessible regions in the SARS-CoV-2 full genome. **Top**: the predicted accessibilities of window size 4 (the blue curve) and 15 (the purple curve) within the 9 regions. **Middle**: accessibilities with window sizes from 1 to 20. The region [25700, 25720] in ORF3a is highly accessible, with an accessibility of greater than 0.9. **Bottom**: the relative position of each accessible region in the full-genome of SARS-CoV-2. Most of the regions are in the last third of the genome, and 3 out of 9 are in the N protein. Note that ORF1ab can be further divided.

## 4. Discussion

We focus on simplifying and accelerating the stochastic sampling algorithm for a given RNA sequence. Algorithmically, we present a hypergraph framework under which the classical sampling algorithm can be greatly simplified. We further elaborate this sampling framework in three versions: the Non-Saving that recovers the hyperedges in a top-down way, the Full-Saving that saves all hyperedges a priori and avoids recomputing for sampling, and the Lazy-Saving that only recovers and saves hyperedges on demand. We show that LazySampling algorithm, i.e., exact partition function followed by a Lazy-Saving sampling, is faster and can avoid unnecessary calculation. Then we present LinearSampling which combines LazySampling and LinearPartition.

LinearSampling is the first algorithm to run in end-to-end linear-time without imposing constraints on the base pair distance, and is orders of magnitude faster than the widely-used Vienna RNAsubopt. We conclude that

- LinearSampling takes linear runtime and can scale up to long RNA sequence, being the first sampling tool to scale to SARS-CoV-2 without local window constraints;
- It approximates the ensemble distribution well;
- Its sampled structures correlate better with the ground truth structures on a diverse database, as well as the experimentally-guided structure of SARS-CoV-2;
- It can be applied to SARS-CoV-2 to discover regions with high accessibilities, which are potential targets for diagnostics and drug design.

## Supporting Information

**Fig. SI 1.**
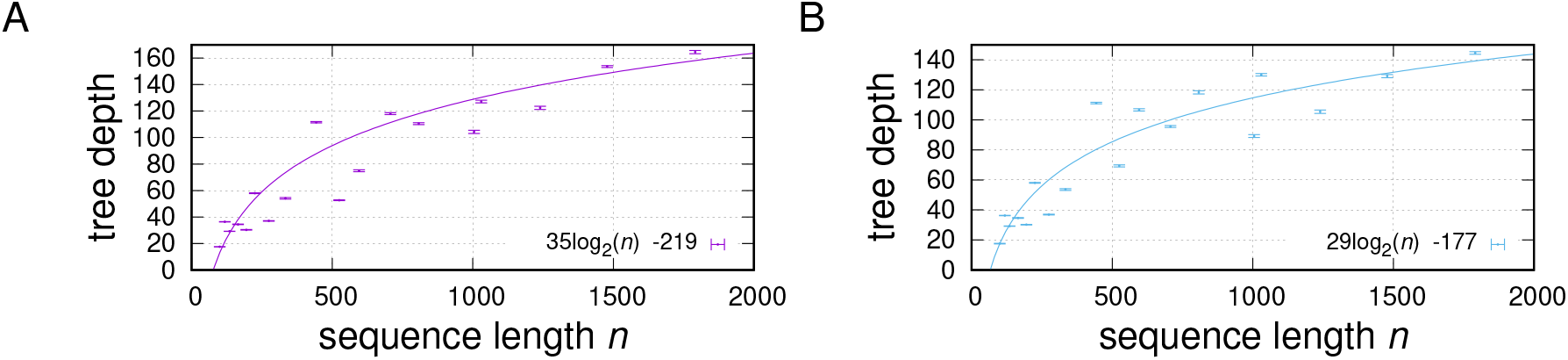
The tree depth of a derivation against sequence length. The horizontal lines show variations across multiple samples (95% confidence intervals). **A**: the tree depth of the structures sampled by Vienna RNAsubopt. **B**: the tree depth of the structures sampled by LinearSampling.

**Fig. SI 2.**
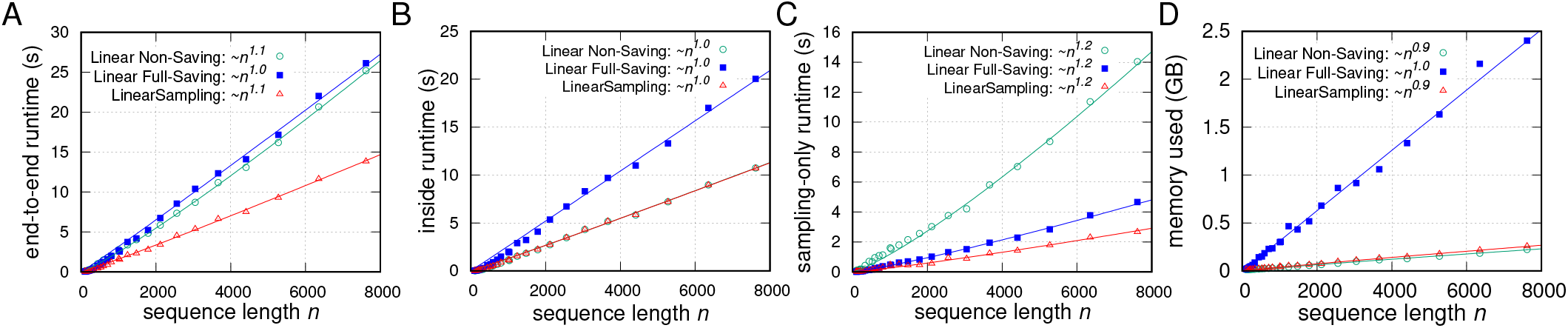
Runtime and memory usage of three sampling strategies, combined with linear partition function, against sequence length *n*. **A**: the end-to-end runtime comparison. **B**: the partition function runtime comparison. **C**: the sampling-only untime comparison. **D**: the memory usage comarison.

**Fig. SI 3.**
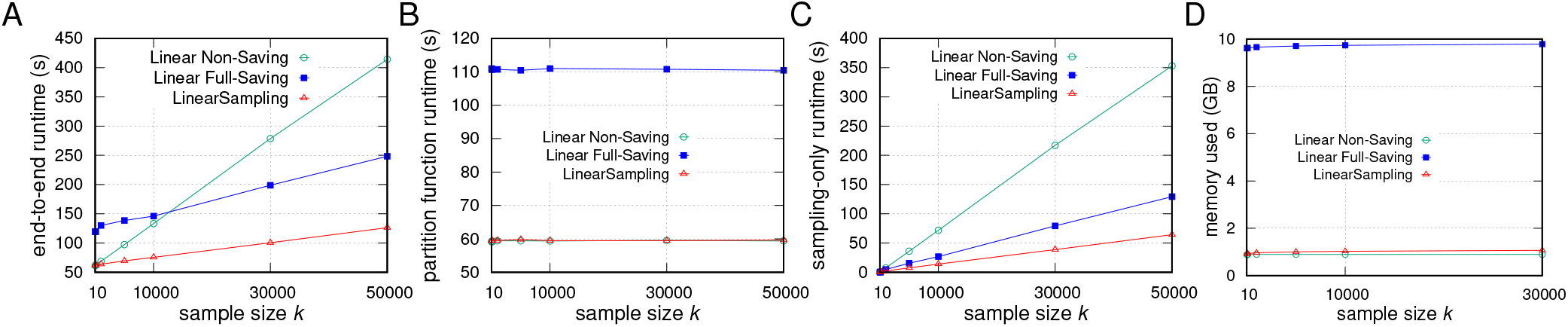
Runtime and memory usage of three sampling strategies, combined with linear partition function, against sample size *k* on the SARS-CoV-2 reference sequence. **A**: the end-to-end runtime comparison. **B**: the partition function runtime comparison. **C**: the sampling-only untime comparison. **D**: the memory usage comarison.

**Fig. SI 4.**
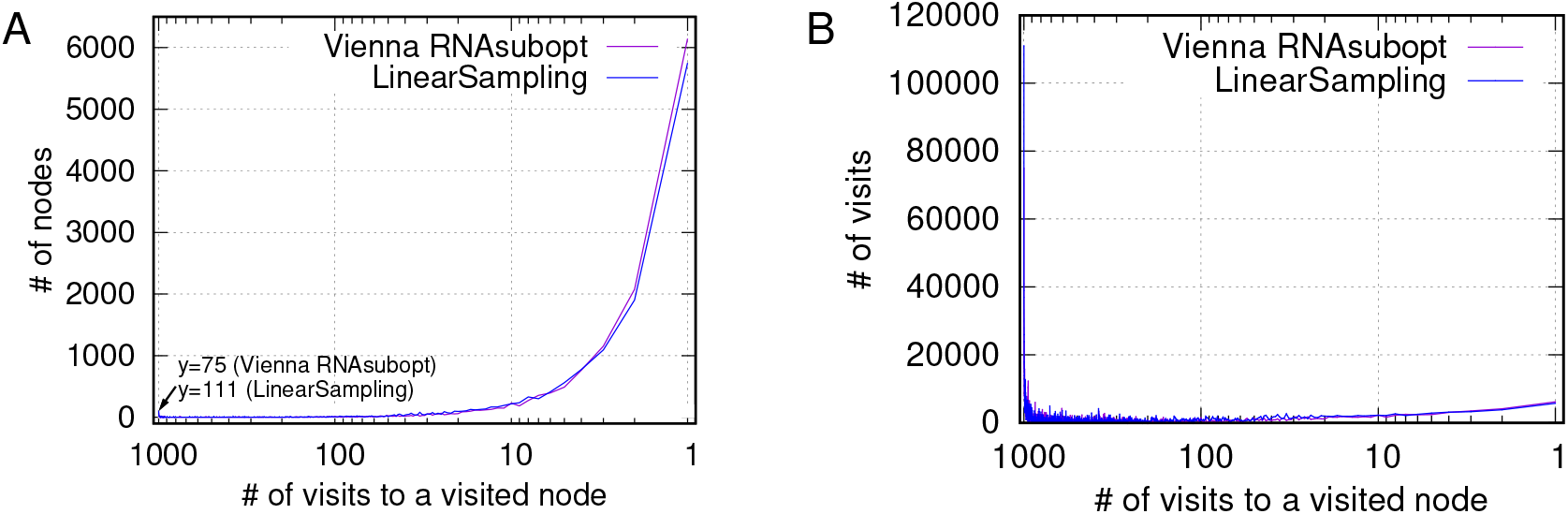
Most of the visits are concentrated on a few nodes. **A**: the distribution of number of nodes against number of visits to a visited node, For example, 111 nodes are visited 1000 times in the sampling phase for LinearSampling. **B**: the distribution of number of visits against number of visits to a visited node.

**Fig. SI 5.**
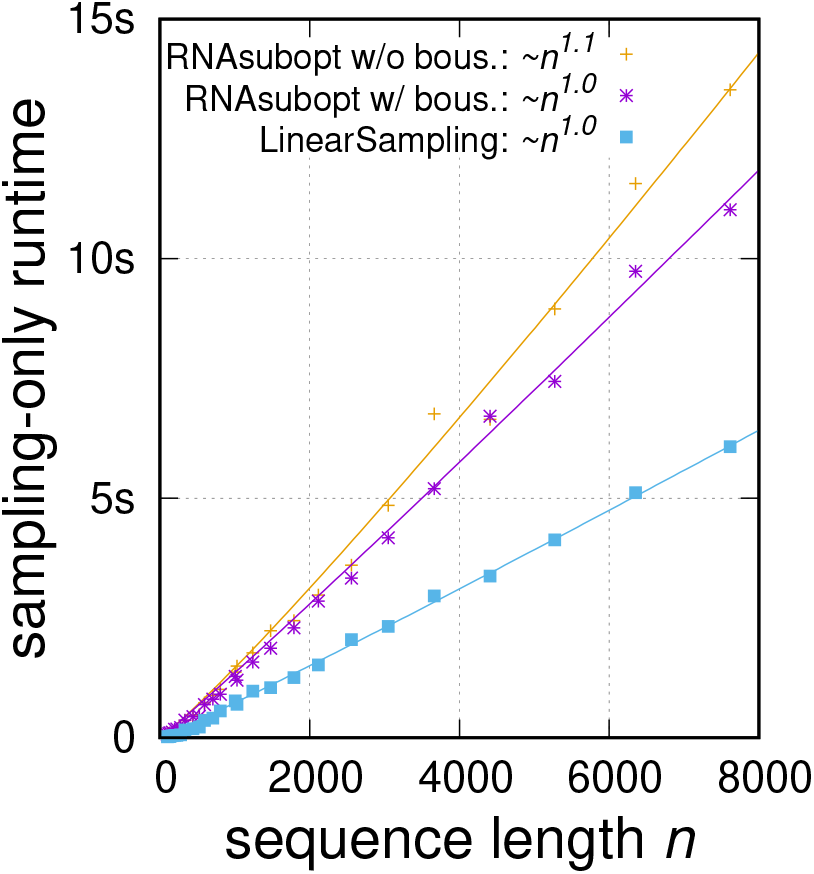
The runtime of Vienna RNAsubopt (with or without the boustrophedon optimization (28)) and LinearSampling, on the RNAcentral dataset up to 8,000 *nt*.

## Footnotes

1 Our notion of “redundant” is unrelated to the one in “non-redundant sampling” (20), which is a variant to output *unique* samples, while standard sampling can sample the same structure more than once.

2 Each empty span **x**_*i,i−1*_ has a special nullary hyperedge 〈**x**_*i,i*−1_, []〉 with no children, and the associated nullary function *f*_0_ is *f*_0_() = “”.

3 For each sample, worst-case: *T*(*n*) = *T*(*n* − 1) + *O*(*n*) = *O*(*n*^2^), and best-case: *T*(*n*) = 2*T*(*n*/2) + *O*(*n*) = *O*(*n* log *n*).

4 Ponty (28) applies the “Boustrophedon” method to reduce the worst-case time also to *O*(*n* log *n*). Our experiments (Fig.SI 5) show that it does further improve the runtime, but only slightly.

5 We point out that RNAstructure (29)’s sampling is similar to our non-saving version except for being non-recursive, i.e., iterative.

6 Ponty (28) analyzes the special case of sampling under the Nussinov model by exploiting the symmetry (though it could have generalized to other systems), and he implemented on the simplified Nussinov-Jacobson (7) rather than the full Turner model (30) as in our work.

7 For each sample, worst-case *T*(*n*) = *T*(*n* − 1) + log *n* = *O*(*n* log *n*); best-case *T*(*n*) = 2*T*(*n*/2) + log *n* = *O*(*n*), similar to “heapify” (31).

8 RNAsubopt local mode does not have a default span size; we choose 70 following the default setting in RNAplfold (36), and 150 since it is the largest default limit in the local folding literature and software.

9 There exist other methods to estimate accessibility, including (a) constrained partition function (forcing each region of interest to be fully unpaired, and compute the fraction of the resulting constrained partition function over the global one), and (b) direct computation (37; 38). But they all run in at least *O*(*n*^3^) time.

